# Community Socioeconomic Disadvantage relates to White Matter Hyperintensity Burden in Mid-to-Late Life Adults

**DOI:** 10.1101/2025.07.01.662674

**Authors:** Anna L. Marsland, Minjie Wu, Mia K. DeCataldo, Howard J. Aizenstein, L. Jinghang, Tamer S. Ibrahim, Stephen B. Manuck, Peter J. Gianaros

## Abstract

**Background:** Residing in communities characterized by socioeconomic disadvantage may confer risk for neurodegenerative brain changes and future neuropathology. Based on prior evidence, this study tested the hypotheses that (1) community-level disadvantage would relate independently of individual-level socioeconomic position to white matter hyperintensities (WMHs), which reflect subclinical brain pathology that may presage later dementia; and (2) this association would be partly explained by blood pressure, cardiometabolic risk, and/or systemic levels of inflammation. These hypotheses were examined among otherwise healthy middle- and older-aged adults without clinical dementia at testing.

**Methods:** Participants were 388 adults aged 40-72 years (53% female; 12% non-White) whose street addresses were entered into the Neighborhood Atlas to compute Area Deprivation Index scores by census block. Participants also underwent high resolution (7 Tesla) brain imaging to assess total WMH volume normalized for intracranial volume, and assessment of blood pressure, cardiometabolic (adiposity, lipids, glucose and insulin), and inflammatory (interleukin-6 and C-reactive protein) risk factors.

**Results:** Linear regression models showed that higher community deprivation on the ADI associated with greater WMH volume, independently of age, sex, years of education and smoking. This association was largely independent of blood pressure, cardiometabolic risk and systemic inflammation.

**Conclusion:** The present novel findings add to growing evidence that community disadvantage relates to preclinical neurodegenerative changes, which may contribute to accelerated brain and cognitive aging. Future work is warranted to better understand pathways that link residential environments to brain health and to identify targets for community and public policy interventions.

## INTRODUCTION

The physical environments and communities in which people reside are recognized as a social determinant of health (Sims et al., 2020), influencing rate of biological aging, independently of chronological age, to shape disparities in population health (Nielsen, Marsland, Hamlat, & Epel, 2024). Living in communities characterized by social and economic disadvantage (e.g., poverty, unemployment, poor housing quality) confers increased risk for cardiometabolic diseases, cerebral vascular disease, neurocognitive decline and dysfunction, incident dementia, and early mortality (Chamberlain et al., 2022; Hunt et al., 2021; Klee, Leist, Veldsman, Ranson, & Llewellyn, 2023; Vassilaki et al., 2022). Community-level disadvantage has also been associated with morphological markers of accelerated brain aging that precede neurocognitive decline and presage risk for dementia in later life. For example, residing in communities characterized by socioeconomic disadvantage has been related to reduced brain tissue volume in the cerebral cortex and the medial temporal lobe (Gianaros et al., 2017; Hunt et al., 2020), decreased cortical surface area and thickness (Hunt et al., 2021; Tan & Tan, 2023), and white matter integrity of midlife adults (Gianaros, Marsland, Sheu, Erickson, & Verstynen, 2013). Moreover, these associations are largely independent of individual-level measures of socioeconomic status (SES), such as indicators of education and income (Gianaros et al., 2017; Hunt et al., 2020). These findings are consistent with postmortem evidence linking community disadvantage to increased risk for Alzheimer’s disease neuropathology (neurofibrillary tangles, cerebral amyloid angiopathy, and neuritic plaque staging)(Hamilton, Matthews, Erskine, Attems, & Thomas, 2021; Powell et al., 2020). The current study extends this work by testing whether community-level disadvantage relates to an established marker of subclinical age-related brain pathology, density of white matter hyperintensities (WMHs).

WMHs are areas of high-intensity signal observed on MRI images that first appear in midlife, increase in prevalence by around 0.4% per year thereafter, and are present in 100% of individuals over age 80 years (de Leeuw et al., 2001; Schmidt et al., 2012). They are thought to reflect damaged brain parenchyma resulting from subclinical cerebral small vessel disease (SVD), although their precise pathophysiology remains unclear and likely reflects diverse etiologies (Debette & Markus, 2010; Kloppenborg, Nederkoorn, Geerlings, & van den Berg, 2014; Prins & Scheltens, 2015). Total WMH burden varies considerably across individuals independently of age and is established as an early biomarker of aging-related brain pathology that antecedes cognitive decline (Debette & Markus, 2010; Tosto, Zimmerman, Carmichael, Brickman, & Alzheimer’s Disease Neuroimaging, 2014), contributes to the pathophysiology of vascular dementia (Gorelick et al., 2011) and risk for Alzheimer’s Disease (Newton et al., 2023) and stroke (Herrmann, Le Masurier, & Ebmeier, 2008). Although not all findings are consistent (Hunt et al., 2020), most studies suggest that individual-level indicators of socioeconomic disadvantage (e.g., education) relate to neurodegenerative brain changes, including WMH burden (Austin et al., 2022; Shaked et al., 2019). However, few studies have examined the independent association of residential area or community-level disadvantage with WMH volume. One exception is the Cardiovascular Health Study, which examined WMH burden and a census derived metric of neighborhood SES (income, education, and occupation) among a population sample of over 3,500 older adults (mean age = 75 years). Cross-sectional results showed no significant association of community-level disadvantage with WMH burden (Rosso et al., 2016); however, living in the lowest income residential areas was associated with a greater increase in WMH volume over a 5-year follow up, a finding that was independent of individual-level SES (Besser et al., 2023). Living in an urban versus a rural setting in India has also been associated with increased WMH load (Aksman, Lynch, Toga, Dey, & Lee, 2023). In the current study, we extend prior work by using a well-validated and broader index of community-level disadvantage that includes physical, social and economic attributes of the residential area. We also focus on a quantitative indicator of burden of WMHs derived from high-field (7 Tesla) brain imaging at a life stage when the prevalence of WMHs begins to accelerate (mean age = 59 years (de Leeuw et al., 2001)) and when preventative intervention may impact trajectories of brain aging and corresponding risk for neuropathology.

Multiple pathways likely contribute to the neurodegeneration that accompanies community-level disadvantage, including biological, behavioral, and social mechanisms. An understanding of these etiological paths is vital to inform prevention and early intervention efforts. Here, we explore three pathogenic mechanisms that may link social and environmental exposures to risk for cerebral SVD and WMHs: age-related increases in blood pressure, cumulative cardiometabolic risk, and systemic levels of inflammatory mediators. Living in a disadvantaged area associates with mid-late-life elevation in cardiometabolic and inflammatory risk factors (e.g., dyslipidemia, elevated blood pressure, adiposity, glucose dysregulation, and systemic levels of inflammatory mediators) and prevalence of their clinical sequelae, cardiovascular, metabolic and chronic inflammatory diseases (Bird et al., 2010; Chamberlain et al., 2022; Diez Roux, 2003; Hu et al., 2021; Keita et al., 2014; Sheets et al., 2020; Vassilaki et al., 2022; Xu, Lawrence, O’Brien, Jackson, & Sandler, 2022). These individual-level factors precede risk for the brain pathology that associates with community-level socioeconomic disadvantage, including cognitive decline, vascular dementia, Alzheimer’s disease, and stroke (Gottesman et al., 2017; Lane et al., 2020; Launer et al., 2000; Thomas et al., 2020), and may contribute to cerebrovascular changes, including lesions to white matter tissue that increase risk for later life neuropathologies (Lane et al., 2020). Recent findings support this possibility, showing that cardiometabolic risk may be a pathway linking community-level socioeconomic disadvantage to reduced brain tissue volume (Gianaros et al., 2017; Gianaros et al., 2023; Hunt et al., 2020). Given evidence that subclinical cardiometabolic risk factors, including blood pressure (Lane et al., 2019), visceral adiposity, systemic inflammation, and fasting glucose levels (Jorgensen et al., 2018), associate with WMH prevalence and predict WMH progression (Lane et al., 2020), we explored the possibility that associations of community-level disadvantage with WMH burden may relate to these possible pathways. Hence, the aims of this study were (1) to test whether community-level disadvantage relates independently of individual-level SES to WMH burden among a sample of relatively healthy mid-late-life community volunteers; and (2) to explore whether this association relates to variability in blood pressure, cumulative cardiometabolic disease risk (a combined index of blood pressure, adiposity, dyslipidemia, and glucose regulation), and/or systemic levels of inflammation (IL-6 and CRP). Community-level disadvantage was assessed using the Area Deprivation Index (ADI), an atlas that ranks census blocks by income, education, employment, and housing quality (Zuelsdorff et al., 2020). It was hypothesized that greater community-level disadvantage measured using the ADI would associate independently of individual-level SES with more WMH burden. Also explored was whether higher blood pressure, cardiometabolic risk, and systemic inflammation are possible pathways linking community-level disadvantage to WMH burden.

## METHODS

### Participants

Data were derived from 388 participants who participated in an MRI as part of wave 2 of the Adult Health and Behavior (AHAB) project. AHAB provides a registry of sociodemographic, behavioral, biological and neural correlates of biological aging and cardiometabolic risk. Participants (30-54 years) were in good general health when recruited via mass-mail solicitation from communities of southwestern Pennsylvania during two time periods (AHAB-I: 2001-2005; AHAB-II: 2008-2011). At enrollment, study exclusions included a reported history of atherosclerotic cardiovascular disease, chronic kidney or liver disease, cancer treatment in the preceding year, major neurological disorders, psychotic illness, systolic/diastolic blood pressure ≥180/110 mmHg (AHAB 1) or ≥160/100 mmHg (AHAB 2), physical incapacities precluding performance of lab tests, pregnancy, and use of insulin, psychotropic, antiarrhythmic, and prescription weight-loss medications ((Natale et al., 2023); See supplement for detail regarding recruitment and eligibility). The present study used data collected in a second wave of AHAB data collection that recruited from both initial cohorts and took place between 2017 and 2024 (12-22 years following initial participation). A total of n = 706 were recruited to participate in this follow-up; however, the MRI sub study was added to the AHAB-1 Wave 2 protocol after the start of the study, so a total of 699 Wave 2 participants were invited to participate in MRI scanning. Neuroimaging data were acquired from n = 388 of the 699, which constitutes the final analytic sample (See supplemental Figure 1 for reasons for missing scan data). When compared with participants who were not scanned (see Table 1), the analytic sample was significantly younger, more likely to endorse being White, reported more years of education, and showed higher cognitive function and lower levels of cardiometabolic risk (See Table 1). They also tended to live in areas characterized by less deprivation on the ADI. Scores on the Montreal Cognitive Assessment (MoCA) administer at wave 2 suggest that average cognitive function of the scanned sample fell in the normal range (mean = 27.1/30; Ratcliffe et al., 2024) All participants denied being diagnosed with dementia. Adjudications were not performed, so clinical determinations regarding mild cognitive impairment could not be made. All Participants provided informed consent, and all procedures were approved by the University of Pittsburgh Institutional Review Board.

**Table 1:** Participant characteristics (mean(standard deviation) or number(percentage) and comparison of the current sample (n = 388) with those who did not complete the MRI (n = 311)

|  | **MRI Not Complete**<br>**(N=311)** | **MRI Complete**<br>**(N=388)** | **Overall**<br>**(N=699)** | **Test Statistic** |
| --- | --- | --- | --- | --- |
| **Age (years)** | 61.3 (6.92) | 58.4 (7.40) | 59.7 (7.33) | t = 5.395*** |
| **Sex at Birth (Male)** | 136 (43.7%) | 181 (46.6%) | 317 (45.4%) | X<sup>2</sup> = 0.482 |
| **Racial and Ethnic Identity** |  |  |  | Fisher Exact* |
| White | 255 (82.0%) | 341 (87.9%) | 596 (85.3%) |  |
| Black | 53 (17.0%) | 39 (10.1%) | 92 (13.2%) |  |
| Asian | 2 (0.6%) | 5 (1.3%) | 7 (1.0%) |  |
| Other | 1 (0.3%) | 3 (0.7%) | 4 (0.6%) |  |
| **Smoking (former/current)** | 116 (37.3%) | 156 (40.2%) | 272 (38.9%) | t = -0.162 |
| **Smoking (Current)** | 27 (8.7%) | 54 (13.9%) | 81 (11.6%) | X<sup>2</sup> = 3.088 |
| **Education (years)** | 16.2 (2.60) | 17.2 (3.10) | 16.7 (2.93) | t = -4.611** |
| **ADI (Percentile)** | 56.7 (24.6) | 52.7 (23.1) | 54.5 (23.8) | t = 2.135* |
| **MoCA** | 26.2 (2.62) | 27.1 (2.40) | 26.7 (2.54) | t = -4.594*** |
| **WMH ICV (mm<sup>3</sup>)** |  | 0.0016(0.0017) |  |  |
| **CMR (Z-units)** | 0.0756 (0.657) | -0.0732 (0.629) | -0.0108 (0.644) | t = 2.881*** |
| **Body Mass Index (kg/m<sup>2</sup>)** | 29.0 (6.58) | 27.8 (5.49) | 28.3 (6.02) | t = 2.432* |
| **Systolic BP (mmHg)** | 127 (15.4) | 124 (15.5) | 126 (15.5) | t = 2.301* |
| **Diastolic BP (mmHg)** | 77.6 (9.75) | 78.1 (9.66) | 77.9 (9.70) | t = -0.672 |
| **IL-6 (pg/mL)** | 2.82 (5.55) | 2.41 (3.54) | 2.58 (4.50) | t = 1.092 |
| **CRP (mg/L)** | 3.21 (8.29) | 2.29 (3.51) | 2.67 (6.02) | t = 1.737 |

- P<.05; \*\* p < .01; \*\*\* p<.001. ADI -Area Deprivation Index; WMH ICT – white matter hyperintensity adjusted for intracranial volume; CMR – continuous cardiometabolic risk; BP- blood pressure; IL-6 – interleukin-6; CRP – C-Reactive Protein; t scores for IL-6 and CRP were calculated using logged values.

### Procedure

Data collected at the first (sociodemographic information, medical history and medication use, blood pressure, and a fasting morning blood draw) and the fourth AHAB wave 2 visit (brain imaging session) were used in the current analyses. Visit 1 was rescheduled if the participant reported symptoms of acute illness or infection, or having had a vaccination in the prior 2 weeks.

### Measures

#### Area Deprivation Index (ADI; (Zuelsdorff et al., 2020)

The ADI provides an estimate of socioeconomic disadvantage of U.S. neighborhoods at the Census block-group level. Higher socioeconomic disadvantage as assessed with this measure has been shown to relate to lower hippocampal and total brain tissue volume(Hunt et al., 2020), Alzheimer disease neuropathology (Powell et al., 2020), and risk for dementia (Chamberlain et al., 2022). The ADI is a composite of 17 area-level US Census metrics assessing education, employment, housing-quality, and poverty measures drawn from the 2013 American Community Survey data (Kind & Buckingham, 2018). Data are publicly available for download on the University of Wisconsin School of Medicine and Public Health’s Neighborhood Atlas website (www.neighborhoodatlas.medicine.wisc.edu). Participants’ street addresses at wave 2 were entered into the Neighborhood Atlas (Version 4) to download ADI values, which are national percentile rankings at the block level (1 = lowest level of neighborhood disadvantage in the US to 100 = highest level of neighborhood disadvantage).

#### White Matter Hyperintensity

To measure total WMH volume, 7-Tesla T2-weighted Fluid Attenuated Inverse Recovery (FLAIR) MR images were obtained using a Siemens 7 T MAGNETOM system equipped with the 1^st^ and 2^nd^ generation Tic Tac Toe (Tac G1 and Tac G2) RF coil systems (Sajewski et al., 2025; Santini et al., 2018; Santini et al., 2021). FLAIR images had a resolution of 0.75 x 0.75x1.5mm and an acquisition matrix of size 240 x 320 x 80. To quantify WMH volume on the FLAIR images, we employed a pretrained deep learning based automatic segmentation model that specializes in WMH delineation across magnetic field strengths and various MR artifacts (Li et al., 2024). Additionally, intracranial volume (ICV) estimations were derived using Freesurfer’s synthstrip toolbox (Hoopes, Mora, Dalca, Fischl, & Hoffmann, 2022).

#### Blood Pressure (BP)

BP was recorded as the average of two consecutive readings of SBP and DBP obtained in a seated position. Mean arterial pressure (MAP) was calculated ((SBP-DBP/3) +DBP). MAP was explored separately from cardiometabolic risk in view of cumulative evidence establishing a unique association of hypertension with cerebral small vessel disease in middle and late adulthood (Hainsworth et al., 2024).

#### Cardiometabolic Risk (CMR)

CMR is known to associate with community disadvantage and metrics of brain aging (Gianaros et al., 2017; Hunt et al., 2020). As in our prior work (Halder et al., 2014), a composite index of CMR modeled on the metabolic syndrome was computed by averaging standardized (z-scored) measures of body mass index, waist circumference (measured at the level of the umbilicus at end-expiration), SBP, and DBP as well as fasting high-density lipoproteins (reverse coded), triglycerides, glucose, and insulin.

#### Inflammatory Index

Serum samples were stored in an ultra-cold freezer and assayed in batches to assess levels of IL-6 and CRP. IL-6 was measured using a high-sensitivity quantitative sandwich enzyme immunoassay kit (range of assay 0.156-10 pg/ml; R&D Systems) following the manufacturer’s directions. Serum CRP levels were measured using a particle-enhanced immunonephelometric assay with the BNII nephelometer from Dade Behring (Newark, DE). The standard range of this assay is 0.175–1,000 mg/L. Intra- and inter-plate coefficients of variation (CVs) for IL-6 and CRP were < 10%. An inflammatory index was calculated by standardizing and averaging the two measures.

#### Standard Covariates

*A priori* covariates included participants self-reported age (in years), sex at birth (coded as male = 1; female = 0), years of education as an index of individual-level SES, and smoking status (coded as current smoker = 1 vs. former- or nonsmoker = 0). We also included age squared as a standard covariate in all models to account for the non-linear relationship between age and prevalence of WMHs (Schmidt et al., 2012).

### Statistical Analyses

All analyses were conducted in R (Version 4.4.1). First, the distributions and number of outliers were assessed. Variables that were skewed or kurtotic were log transformed, which included IL-6, CRP, triglycerides, and glucose. WMH values were first normalized by intracranial volumes and also log-transformed for analysis. Next, descriptive bivariate correlations between ADI values, years of education, CMR (and its components), inflammation, and WMH adjusted for intracranial volume were examined (See Supplemental Table 1). Fixed-effect ordinary least squares linear regression models were used to test the association between neighborhood disadvantage and WMH adjusted for intracranial volume, with age, sex at birth, years of education, and smoking status included as standard covariates. Missing data were imputed in these models using full information maximum likelihood. To explore the possibility that CMR, MAP, and inflammation contributed to the relationship of ADI values with WMH, we added each of them separately to the initial regression model. Finally, we explored the possibility that biological sex at birth moderated the association of ADI values with WMH volume. There were no more than 2 participants living in a single census block so nested analyses were not performed.

## RESULTS

Table 1 contains descriptive statistics for the sample (n = 388; mean age 58 (SD = 7.4; range 40-72) years; 53.4% identified as female at birth; 12.1% identified as Black, Asian, Indigenous or Multiracial and 87.9% as White). On average, the sample was well educated with 17.2 (range 10-31) years of schooling. Scores on the ADI spanned the full range of deprivation in the US, with census-tracts based on participants residential address ranging from the 4^th^ to the 100^th^ percentile (mean = 53rd percentile). Bivariate correlations are presented in Supplementary Table S1. Higher ICV-adjusted WMH volume was associated with living in more deprived neighborhoods on the ADI (r = .11, p = .04; See Figure 1). In addition, adjusted WMH significantly associated with older age (See Figure 2), and higher mean arterial (See Figure 3), systolic and diastolic BP, higher glucose levels, waist circumference, continuous CMR, the inflammatory composite, and circulating levels of IL-6 (p’s < .05; see supplementary Table 1). Higher levels of community-level disadvantage on the ADI associated significantly with fewer years of education, higher mean arterial and DBP, higher waist circumference, BMI, continuous CMR, circulating levels of insulin, IL-6 and CRP, and the combined inflammatory score (p’s<.05).

**Figure 1.**
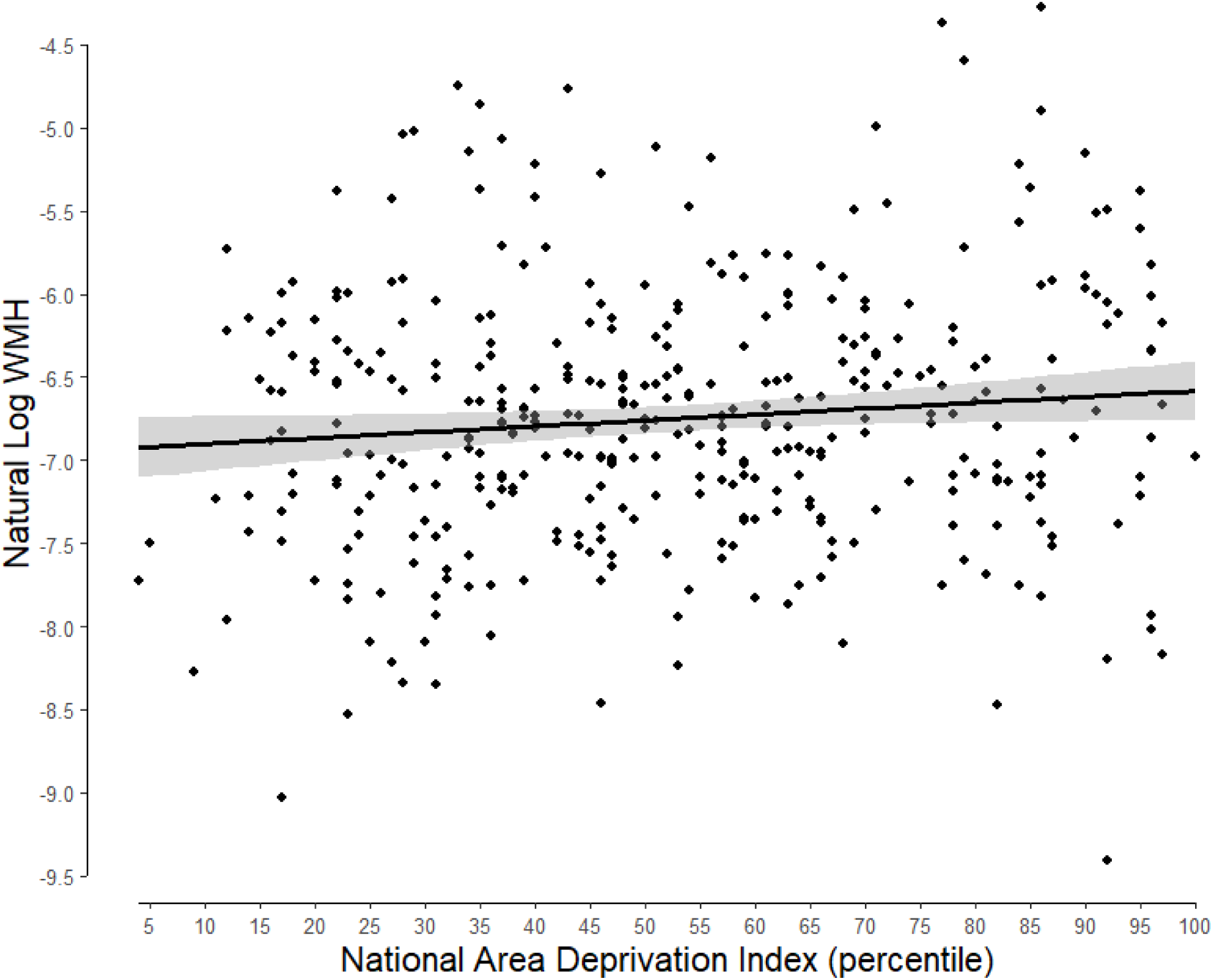
Scatterplot showing the relationship between Area Deprivation Index percentile and intracranial volume-adjusted white matter hyperintensities.

**Figure 2.**
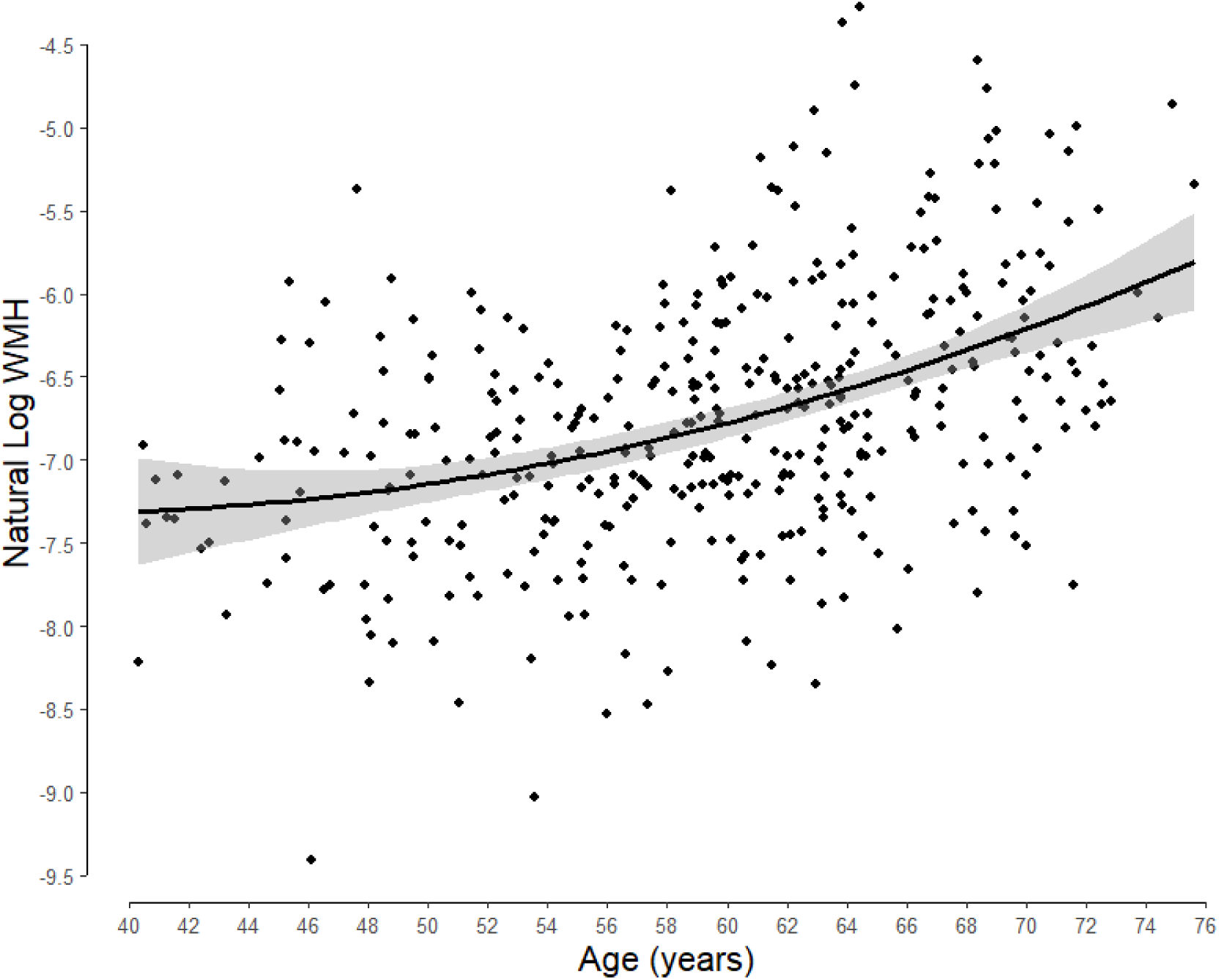
Scatterplot showing the relationship between age and intracranial volume-adjusted white matter hyperintensities.

**Figure 3.**
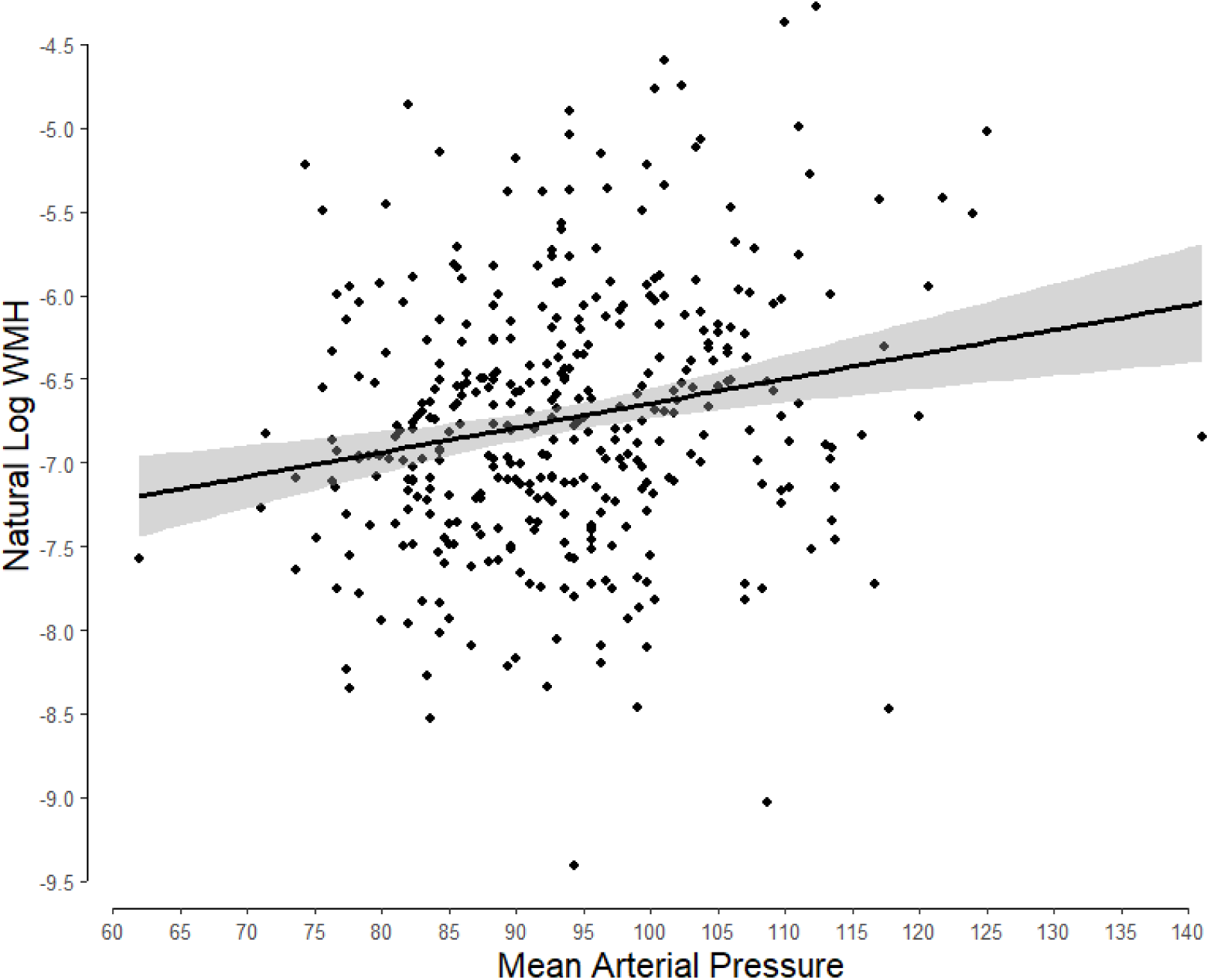
Scatterplot showing the relationship between mean aretrial pressure and intracranial volume-adjusted white matter hyperintensities.

Results of the primary regression model are displayed in Table 2. ADI percentiles associated with ICV-adjusted WMH volume (beta = 0.10, p = .03) independently of age^2^, sex at birth, years of education, and smoking status. We next examined the impact of separately adding CMR, the inflammatory index, and MAP to the primary regression model (Table 3). The observed association of ADI percentile with ICV-adjusted WMH density remained significant in models that included CMR (β = 0.10, p = .043) and the inflammatory index (β = 0.10, p = .045), with no independent association of either of these factors with WMH. There was a slight attenuation of the relationship of ADI percentile with ICV-adjusted WMH in the model that included MAP (β = 0.09, p = .069). In this model, there was also an independent relationship of MAP with density of ICV-adjusted WMH (β = .14, p = .004). Exploratory mediational analyses showed no significant indirect effect of ADI percentiles on ICV-adjusted WMH density through MAP (Indirect effect coefficient = 0.012; 95% confident intervals: -0.19 to 0.87; p = 0.15). Finally, results of models that included the interaction of biological sex at birth and ADI percentiles showed no significant sex difference in the association of neighborhood disadvantage with WMH density (see supplementary Table2).

**Table 2.** Results of regression model predicting white matter hyperintensities volume adjusted for intracranial volume.

| | $\beta$ | 95% CIs | p |
| --- | --- | --- | --- |
| ADI | 0.10 | 0.01 to 0.20 | **0.03** |
| Age<sup>2</sup> | 0.45 | 0.36 to 0.54 | **<0.001** |
| Sex at Birth | -0.13 | -0.23 to -0.04 | **0.004** |
| Years of Education | -0.00 | -0.10 to 0.10 | 0.99 |
| Smoking Status | -0.00 | -0.09 to 0.09 | 0.95 |

ADI – Area Deprivation Index; CI – confidence interval

**Table 3.**
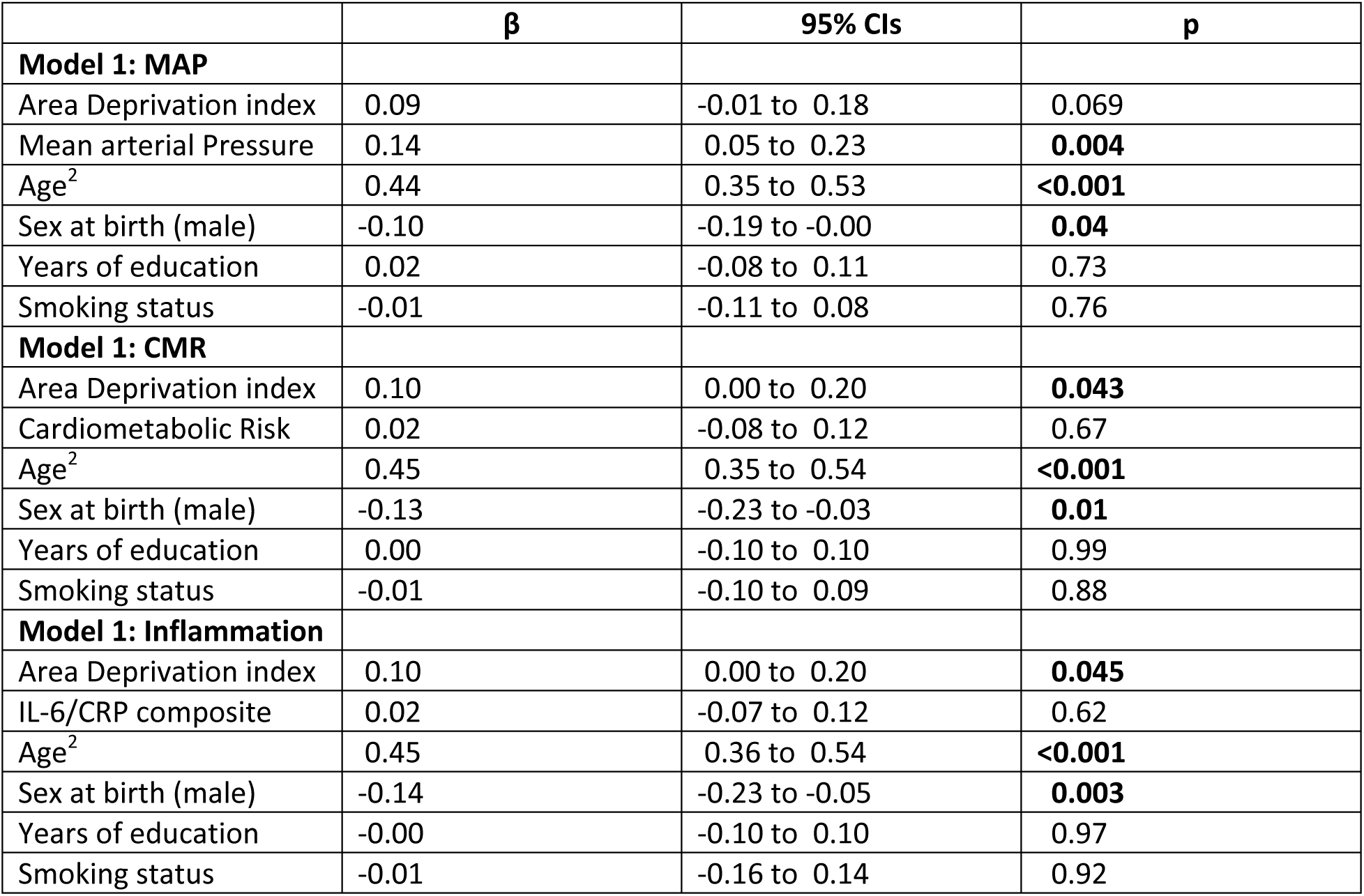
Results of separate models adding mean arterial pressure (MAP), continous cardiometabolci risk (CMR) and the inflammatory composite.

| | $\beta$ | 95% CIs | p |
| --- | --- | --- | --- |
| **Model 1: MAP** |  |  |  |
| Area Deprivation index | 0.09 | -0.01 to 0.18 | 0.069 |
| Mean arterial Pressure | 0.14 | 0.05 to 0.23 | **0.004** |
| Age<sup>2</sup> | 0.44 | 0.35 to 0.53 | **<0.001** |
| Sex at birth (male) | -0.10 | -0.19 to -0.00 | **0.04** |
| Years of education | 0.02 | -0.08 to 0.11 | 0.73 |
| Smoking status | -0.01 | -0.11 to 0.08 | 0.76 |
| **Model 1: CMR** |  |  |  |
| Area Deprivation index | 0.10 | 0.00 to 0.20 | **0.043** |
| Cardiometabolic Risk | 0.02 | -0.08 to 0.12 | 0.67 |
| Age<sup>2</sup> | 0.45 | 0.35 to 0.54 | **<0.001** |
| Sex at birth (male) | -0.13 | -0.23 to -0.03 | **0.01** |
| Years of education | 0.00 | -0.10 to 0.10 | 0.99 |
| Smoking status | -0.01 | -0.10 to 0.09 | 0.88 |
| **Model 1: Inflammation** |  |  |  |
| Area Deprivation index | 0.10 | 0.00 to 0.20 | **0.045** |
| IL-6/CRP composite | 0.02 | -0.07 to 0.12 | 0.62 |
| Age<sup>2</sup> | 0.45 | 0.36 to 0.54 | **<0.001** |
| Sex at birth (male) | -0.14 | -0.23 to -0.05 | **0.003** |
| Years of education | -0.00 | -0.10 to 0.10 | 0.97 |
| Smoking status | -0.01 | -0.16 to 0.14 | 0.92 |

## DISCUSSION

The residential environments in which people live, work, play, and age affect their health, functioning and quality of life (Sims et al., 2020). The present study examined whether a well-validated socioeconomic index of residential area (community-level) conditions relates to WMH burden, a marker of brain pathology that presages risk for cognitive decline, dementia (Debette & Markus, 2010; Gorelick et al., 2011; Kloppenborg et al., 2014; Newton et al., 2023; Prins & Scheltens, 2015; Tosto et al., 2014) and stroke (Herrmann et al., 2008). Among a relatively healthy sample of 387 mid-to-late life volunteers, findings showed that residing in a more disadvantaged community associated with greater WMH burden, a relationship that was independent of individual SES, as indexed by years of education. These findings contribute to evidence that living in disadvantaged areas relates to preclinical neurodegeneration, as previously indexed by cortical thinning, lower global brain and hippocampal volume (Gianaros et al., 2017; Hunt et al., 2020; Hunt et al., 2021), Alzheimer’s Disease neuropathology (Powell et al., 2020), and accelerated cognitive aging (Kuchibhatla et al., 2020; Rosso et al., 2016; Vassilaki et al., 2022; Zuelsdorff et al., 2020). Two prior studies have examined the association of community disadvantage with WMC burden. The first examined a later life sample (mean age 75 years) and showed no significant cross-sectional relationship of a census-derived index of area-level SES (block level metrics of wealth, education, and occupation) with WMH volume (Rosso et al., 2016), but that living in the lowest income areas associated with a larger increase in WMH volume over a 5-year follow-up than living in more advantaged areas (Besser et al., 2023). The second study examined data on 19,638 individuals (mean age 55 years) from the UK Biobank. Here, area-level SES was assessed with the Townsend deprivation index, which includes levels of unemployment, car and home ownership, and overcrowding. Results showed an association of living in a high deprivation area with greater WMH burden, but only among individuals of lower individual-level SES (Tan & Tan, 2023). The current findings extend prior work by (1) examining a broader index of area-level conditions, including physical, economic, and social community attributes, (2) employing a more sensitive assessment of WMH burden derived from the higher spatial resolution afforded by 7 Tesla brain imaging, and (3) assessing individuals at a life stage when the prevalence of WMHs begins to accelerate (mean age = 59 years (de Leeuw et al., 2001). Our results provide further support for the relationship of community attributes with preclinical neurodegeneration, supporting the value of investigating the specific aspects of area and environmental exposures and timing of exposures across the life course that precipitate WMH development in order to maximize prevention and treatment success.

Higher individual-level SES as indexed by income and education is widely believed to protect the brain and preserve cognitive function in later life, possibly as a function of increased cognitive reserve (Jefferson et al., 2011). However, results of studies examining the relation of individual-level SES to WMH burden and other preclinical markers of brain aging are mixed. The current findings are consistent with studies that show no clear relation of education with WMH burden (Eng et al., 2021; Rodriguez et al., 2023) or other preclinical markers of brain aging, including regional cortical thickness and gray matter volume, among non-clinical samples (Fotenos, Mintun, Snyder, Morris, & Buckner, 2008; Liu et al., 2012; Mungas et al., 2018; Seo et al., 2011; Walhovd et al., 2022). Other studies do show an association of less education with accelerated age-related neurodegeneration (Kim et al., 2015; Shaked et al., 2019). In one of the only studies to examine WMH burden, results show an inverse association of individual-level SES as indexed by education and poverty status, with WMH burden, but only among African Americans (Waldstein et al., 2017). These results raise the possibility that environmental exposures contribute to patterns of relationships, particularly given that race, individual SES and area-level exposures are often confounded in the United States (Williams, Mohammed, Leavell, & Collins, 2012), making it difficult to untangle their relative contributions. In the current study higher education was related to ADI values (r = -0.33), reflecting living in a more advantaged neighborhood; however, there was no significant association of education with WMH burden, suggesting that aspects of the communities in which individuals reside contribute more to WHM load than individual-level disadvantage.

Living in communities characterized by social, physical and economic disadvantage increases exposure to unfavorable life circumstances and environmental factors that may contribute to the development of WMHs and negatively affect brain health (Adler & Stewart, 2010). In the current study, we explored three possible pathogenic mechanisms: age-related increases in blood pressure, cumulative cardiometabolic risk, and systemic levels of inflammation. Consistent with existing literature (Bird et al., 2010; Chamberlain et al., 2022; Diez Roux, 2003; Hu et al., 2021; Keita et al., 2014; Sheets et al., 2020; Vassilaki et al., 2022; Xu et al., 2022), residing in a more disadvantaged community was associated with elevated blood pressure, increased cardiometabolic risk, and higher levels of systemic inflammation. However, the association of our measure of community-level disadvantage (ADI values) with WMH burden was largely independent of these risk pathways. Consistent with evidence that higher blood pressure contributes to the pathogenesis of WMH lesions (Hainsworth, Markus, & Schneider, 2024; Lane et al., 2020; Lane et al., 2019), we did observe a positive association of blood pressure with WMH burden that was independent of ADI values. Other factors that may contribute to the observed association of ADI values with WMH burden include historical structural influences and inequitable policies that have resulted in neighborhoods and residential areas that differ in access to resources (e.g., places to exercise, healthy food and other amenities that support a healthy lifestyle), employment opportunities and healthcare, and exposure to hazards (e.g., pollution, toxic materials, levels of crime, threats to safety, and psychological stress) (Diez Roux & Mair, 2010). It is widely acknowledged that these physical and social community attributes shape the physical and mental health at the population level, contributing to observed geographic health disparities (Egede, Walker, & Williams, 2024). We previously reported that exposure to toxic pollution, elevated homicide rates, lack of green space, fewer employment opportunities, and limited area resources associate with decreased cortical tissue volume, a morphological marker of accelerated brain aging (Gianaros et al., 2023). Furthermore, these associations were independent of and in addition to a census-based composite of area-level socioeconomic disadvantage. Other work suggests that psychosocial factors, including chronic stress, social isolation and psychopathology, may contribute to the accelerated brain aging that accompanies living in disadvantaged neighborhoods (Livingston et al., 2020). Future work should examine the possibility that lifestyle behaviors, psychosocial factors, and physical exposures contribute to the association of neighborhood and area-level living conditions with burden of WMHs.

The current study is limited by its cross-sectional design, which prevents causal interpretation. Recent longitudinal evidence shows that higher levels of community-level deprivation on the ADI predict accelerated preclinical neurodegeneration and cognitive decline, and increased risk for progression to dementia among community dwelling mid-late life adults who were cognitively unimpaired at baseline (Hunt et al., 2021; Vassilaki et al., 2022). The current findings raise the possibility that WMHs may contribute to this functional decline. However, alternative interpretations of the current cross-sectional findings should also be considered. For example, it is possible that neurodegeneration and associated cognitive decline precede movement into disadvantaged residential areas. The assessment of ADI at one timepoint in mid-late adulthood also precludes the consideration of duration and timing of exposures. Recent findings suggest that risk for premature mortality is greatest among individuals who reside in persistently disadvantaged neighborhoods from early to middle adulthood, with low neighborhood disadvantage in childhood, young adulthood and mid-life all associating with elevated risk (Lawrence et al., 2024). It is also possible that environmental exposures in early life (e.g., undernutrition) program the developing brain and shift lifelong trajectories of neurocognitive health in ways that may impact socioeconomic circumstances in later life (Jones et al., 2019).

The current study is also limited by the enrollment of predominantly non-Hispanic White and relatively advantaged volunteers. Although the sample included participants from the full range of neighborhood disadvantage, as assessed by the ADI, the low number of participants falling in the most deprived ADI rankings suggests individuals residing in lower socioeconomic circumstances are underrepresented. It is therefore possible that we underestimated the possible impact of community-level disadvantage. The study sample also limits generalizability to those living outside a mostly urban setting and was not statistically powered to consider the intersection of race and ethnicity with socioeconomic and community factors. In this regard, future research needs to address the extent that racist historical policies have impacted environmental exposures, psychosocial factors and access to resources that may impact brain aging (Pohl et al., 2021). Future longitudinal work is warranted examining the impact of a broad range of residential area exposures across the lifespan and at critical periods for brain development on brain aging in later life. This work should extend to include the assessment of cognitive changes that accompany the development of WMHs.

In conclusion, the present findings contribute to a burgeoning literature suggesting that the environments in which individuals reside are impactful for brain health. Here, we show an association of residential area-level disadvantage with a subclinical marker of age-related brain pathology known to antecede cognitive decline and predict risk for dementia and stroke among a sample of community dwelling mid-late-life adults. These findings extend prior work by examining a broader index of community-level disadvantage that considers physical and social attributes of residential environments, as well as a quantitative measure of WMH derived from high resolution brain images. Future work is warranted to better understand pathways that link residential community exposures to brain health and to identify targets for population and public policy interventions.

## Acknowledgements

We thank all the participants and staff who contributed to the AHAB study.

## REFERENCES

Adler, N. E., & Stewart, J. (2010). Health disparities across the lifespan: meaning, methods, and mechanisms. Annals of the New York Academy of Sciences, 1186, 5–23. doi:10.1111/j.1749-6632.2009.05337.x

Aksman, L., Lynch, K., Toga, A., Dey, A. B., & Lee, J. (2023). Investigating the factors that explain white matter hyperintensity load in older Indians. Brain Commun, 5(1), fcad008. doi:10.1093/braincomms/fcad008

Austin, T. R., Nasrallah, I. M., Erus, G., Desiderio, L. M., Chen, L. Y., Greenland, P., . . . Heckbert, S. R. (2022). Association of Brain Volumes and White Matter Injury With Race, Ethnicity, and Cardiovascular Risk Factors: The Multi-Ethnic Study of Atherosclerosis. J Am Heart Assoc, 11(7), e023159. doi:10.1161/JAHA.121.023159

Besser, L. M., Lovasi, G. S., Zambrano, J. J., Camacho, S., Dhanekula, D., Michael, Y. L., . . . Longstreth, W. T. (2023). Neighborhood greenspace and neighborhood income associated with white matter grade worsening: Cardiovascular Health Study. Alzheimers Dement (Amst), 15(4), e12484. doi:10.1002/dad2.12484

Bird, C. E., Seeman, T., Escarce, J. J., Basurto-Dávila, R., Finch, B. K., Dubowitz, T., . . . Lurie, N. (2010). Neighbourhood socioeconomic status and biological ’wear and tear’ in a nationally representative sample of US adults. Journal of Epidemiology and Community Health, 64(10), 860–865. doi:10.1136/jech.2008.084814

Chamberlain, A. M., St Sauver, J. L., Finney Rutten, L. J., Fan, C., Jacobson, D. J., Wilson, P. M., . . . Rocca, W. A. (2022). Associations of Neighborhood Socioeconomic Disadvantage With Chronic Conditions by Age, Sex, Race, and Ethnicity in a Population-Based Cohort. Mayo Clinic Proceedings, 97(1), 57–67. doi:10.1016/j.mayocp.2021.09.006

de Leeuw, F. E., de Groot, J. C., Achten, E., Oudkerk, M., Ramos, L. M., Heijboer, R., . . . Breteler, M. M. (2001). Prevalence of cerebral white matter lesions in elderly people: a population based magnetic resonance imaging study. The Rotterdam Scan Study. Journal of Neurology, Neurosurgery and Psychiatry, 70(1), 9–14. doi:10.1136/jnnp.70.1.9

Debette, S., & Markus, H. S. (2010). The clinical importance of white matter hyperintensities on brain magnetic resonance imaging: systematic review and meta-analysis. BMJ, 341, c3666. doi:10.1136/bmj.c3666

Diez Roux, A. V. (2003). Residential environments and cardiovascular risk. Journal of Urban Health, 80(4), 569–589. doi:10.1093/jurban/jtg065

Diez Roux, A. V., & Mair, C. (2010). Neighborhoods and health. Annals of the New York Academy of Sciences, 1186, 125–145. doi:10.1111/j.1749-6632.2009.05333.x

Egede, L. E., Walker, R. J., & Williams, J. S. (2024). Addressing Structural Inequalities, Structural Racism, and Social Determinants of Health: a Vision for the Future. Journal of General Internal Medicine, 39(3), 487–491. doi:10.1007/s11606-023-08426-7

Eng, C. W., Gilsanz, P., Fletcher, E. M., Kornak, J., George, K. M., DeCarli, C. S., . . . Whitmer, R. A. (2021). Education, white matter hyperintnsities, and cognition in diverse cohorts of older adults. Alzheimers & Dementia, 17, 1–3.

Fotenos, A. F., Mintun, M. A., Snyder, A. Z., Morris, J. C., & Buckner, R. L. (2008). Brain volume decline in aging: evidence for a relation between socioeconomic status, preclinical Alzheimer disease, and reserve. Archives of Neurology, 65(1), 113–120. doi:10.1001/archneurol.2007.27

Gianaros, P. J., Kuan, D. C., Marsland, A. L., Sheu, L. K., Hackman, D. A., Miller, K. G., & Manuck, S. B. (2017). Community Socioeconomic Disadvantage in Midlife Relates to Cortical Morphology via Neuroendocrine and Cardiometabolic Pathways. Cerebral Cortex, 27(1), 460–473. doi:10.1093/cercor/bhv233

Gianaros, P. J., Marsland, A. L., Sheu, L. K., Erickson, K. I., & Verstynen, T. D. (2013). Inflammatory pathways link socioeconomic inequalities to white matter architecture. Cerebral Cortex, 23(9), 2058–2071. doi:10.1093/cercor/bhs191

Gianaros, P. J., Miller, P. L., Manuck, S. B., Kuan, D. C., Rosso, A. L., Votruba-Drzal, E. E., & Marsland, A. L. (2023). Beyond Neighborhood Disadvantage: Local Resources, Green Space, Pollution, and Crime as Residential Community Correlates of Cardiovascular Risk and Brain Morphology in Midlife Adults. Psychosomatic Medicine, 85(5), 378–388. doi:10.1097/PSY.0000000000001199

Gorelick, P. B., Scuteri, A., Black, S. E., Decarli, C., Greenberg, S. M., Iadecola, C., . . . Anesthesia. (2011). Vascular contributions to cognitive impairment and dementia: a statement for healthcare professionals from the american heart association/american stroke association. Stroke, 42(9), 2672–2713. doi:10.1161/STR.0b013e3182299496

Gottesman, R. F., Albert, M. S., Alonso, A., Coker, L. H., Coresh, J., Davis, S. M., . . . Knopman, D. S. (2017). Associations Between Midlife Vascular Risk Factors and 25-Year Incident Dementia in the Atherosclerosis Risk in Communities (ARIC) Cohort. JAMA Neurol, 74(10), 1246–1254. doi:10.1001/jamaneurol.2017.1658

Hainsworth, A. H., Markus, H. S., & Schneider, J. A. (2024). Cerebral Small Vessel Disease, Hypertension, and Vascular Contributions to Cognitive Impairment and Dementia. Hypertension, 81(1), 75–86. doi:10.1161/HYPERTENSIONAHA.123.19943

Halder, I., Champlin, J., Sheu, L., Goodpaster, B. H., Manuck, S. B., Ferrell, R. E., & Muldoon, M. F. (2014). PPARalpha gene polymorphisms modulate the association between physical activity and cardiometabolic risk. Nutrition, Metabolism, and Cardiovascular Diseases, 24(7), 799–805. doi:10.1016/j.numecd.2014.02.007

Hamilton, C. A., Matthews, F. E., Erskine, D., Attems, J., & Thomas, A. J. (2021). Neurodegenerative brain changes are associated with area deprivation in the United Kingdom: findings from the Brains for Dementia Research study. Acta Neuropathol Commun, 9(1), 198. doi:10.1186/s40478-021-01301-8

Herrmann, L. L., Le Masurier, M., & Ebmeier, K. P. (2008). White matter hyperintensities in late life depression: a systematic review. Journal of Neurology, Neurosurgery and Psychiatry, 79(6), 619–624. doi:10.1136/jnnp.2007.124651

Hoopes, A., Mora, J. S., Dalca, A. V., Fischl, B., & Hoffmann, M. (2022). SynthStrip: skull-stripping for any brain image. Neuroimage, 260, 119474.

Hu, M. D., Lawrence, K. G., Bodkin, M. R., Kwok, R. K., Engel, L. S., & Sandler, D. P. (2021). Neighborhood Deprivation, Obesity, and Diabetes in Residents of the US Gulf Coast. American Journal of Epidemiology, 190(2), 295–304. doi:10.1093/aje/kwaa206

Hunt, J. F. V., Buckingham, W., Kim, A. J., Oh, J., Vogt, N. M., Jonaitis, E. M., . . . Bendlin, B. B. (2020). Association of Neighborhood-Level Disadvantage With Cerebral and Hippocampal Volume. JAMA Neurol, 77(4), 451–460. doi:10.1001/jamaneurol.2019.4501

Hunt, J. F. V., Vogt, N. M., Jonaitis, E. M., Buckingham, W. R., Koscik, R. L., Zuelsdorff, M., . . . Kind, A. J. H. (2021). Association of Neighborhood Context, Cognitive Decline, and Cortical Change in an Unimpaired Cohort. Neurology, 96(20), e2500–e2512. doi:10.1212/WNL.0000000000011918

Jefferson, A. L., Gibbons, L. E., Rentz, D. M., Carvalho, J. O., Manly, J., Bennett, D. A., & Jones, R. N. (2011). A life course model of cognitive activities, socioeconomic status, education, reading ability, and cognition. Journal of the American Geriatrics Society, 59(8), 1403–1411. doi:10.1111/j.1532-5415.2011.03499.x

Jones, N. L., Gilman, S. E., Cheng, T. L., Drury, S. S., Hill, C. V., & Geronimus, A. T. (2019). Life Course Approaches to the Causes of Health Disparities. American Journal of Public Health, 109(S1), S48–S55. doi:10.2105/AJPH.2018.304738

Jorgensen, D. R., Shaaban, C. E., Wiley, C. A., Gianaros, P. J., Mettenburg, J., & Rosano, C. (2018). A population neuroscience approach to the study of cerebral small vessel disease in midlife and late life: an invited review. American Journal of Physiology: Heart and Circulatory Physiology, 314(6), H1117–H1136. doi:10.1152/ajpheart.00535.2017

Keita, A. D., Judd, S. E., Howard, V. J., Carson, A. P., Ard, J. D., & Fernandez, J. R. (2014). Associations of neighborhood area level deprivation with the metabolic syndrome and inflammation among middle- and older-age adults. BMC Public Health, 14, 1319. doi:10.1186/1471-2458-14-1319

Kim, J. P., Seo, S. W., Shin, H. Y., Ye, B. S., Yang, J. J., Kim, C., . . . Guallar, E. (2015). Effects of education on aging-related cortical thinning among cognitively normal individuals. Neurology, 85(9), 806–812. doi:10.1212/WNL.0000000000001884

Kind, A. J. H., & Buckingham, W. R. (2018). Making Neighborhood-Disadvantage Metrics Accessible - The Neighborhood Atlas. New England Journal of Medicine, 378(26), 2456–2458. doi:10.1056/NEJMp1802313

Klee, M., Leist, A. K., Veldsman, M., Ranson, J. M., & Llewellyn, D. J. (2023). Socioeconomic Deprivation, Genetic Risk, and Incident Dementia. American Journal of Preventive Medicine, 64(5), 621–630. doi:10.1016/j.amepre.2023.01.012

Kloppenborg, R. P., Nederkoorn, P. J., Geerlings, M. I., & van den Berg, E. (2014). Presence and progression of white matter hyperintensities and cognition: a meta-analysis. Neurology, 82(23), 2127–2138. doi:10.1212/WNL.0000000000000505

Kuchibhatla, M., Hunter, J. C., Plassman, B. L., Lutz, M. W., Casanova, R., Saldana, S., & Hayden, K. M. (2020). The association between neighborhood socioeconomic status, cardiovascular and cerebrovascular risk factors, and cognitive decline in the Health and Retirement Study (HRS). Aging Ment Health, 24(9), 1479–1486. doi:10.1080/13607863.2019.1594169

Lane, C. A., Barnes, J., Nicholas, J. M., Sudre, C. H., Cash, D. M., Malone, I. B., . . . Schott, J. M. (2020). Associations Between Vascular Risk Across Adulthood and Brain Pathology in Late Life: Evidence From a British Birth Cohort. JAMA Neurol, 77(2), 175–183. doi:10.1001/jamaneurol.2019.3774

Lane, C. A., Barnes, J., Nicholas, J. M., Sudre, C. H., Cash, D. M., Parker, T. D., . . . Schott, J. M. (2019). Associations between blood pressure across adulthood and late-life brain structure and pathology in the neuroscience substudy of the 1946 British birth cohort (Insight 46): an epidemiological study. Lancet Neurology, 18(10), 942–952. doi:10.1016/S1474-4422(19)30228-5

Launer, L. J., Ross, G. W., Petrovitch, H., Masaki, K., Foley, D., White, L. R., & Havlik, R. J. (2000). Midlife blood pressure and dementia: the Honolulu-Asia aging study. Neurobiology of Aging, 21(1), 49–55. doi:10.1016/s0197-4580(00)00096-8

Lawrence, W. R., Kucharska-Newton, A. M., Magnani, J. W., Brewer, L. C., Shiels, M. S., George, K. M., . . . Freedman, N. D. (2024). Neighborhood Socioeconomic Disadvantage Across the Life Course and Premature Mortality. JAMA Netw Open, 7(8), e2426243. doi:10.1001/jamanetworkopen.2024.26243

Li, J., Santini, T., Huang, Y., Mettenburg, J. M., Ibrahima, T. S., Aizensteina, H. J., & Wu, M. (2024). wmh_seg: Transformer based U-Net for Robust and Automatic White Matter Hyperintensity Segmentation across 1.5 T, 3T and 7T. arXiv preprint arXiv:2402.12701.

Liu, Y., Julkunen, V., Paajanen, T., Westman, E., Wahlund, L. O., Aitken, A., . . . AddNeuroMed, C. (2012). Education increases reserve against Alzheimer’s disease--evidence from structural MRI analysis. Neuroradiology, 54(9), 929–938. doi:10.1007/s00234-012-1005-0

Livingston, G., Huntley, J., Sommerlad, A., Ames, D., Ballard, C., Banerjee, S., . . . Mukadam, N. (2020). Dementia prevention, intervention, and care: 2020 report of the Lancet Commission. Lancet, 396(10248), 413–446. doi:10.1016/S0140-6736(20)30367-6

Mungas, D., Gavett, B., Fletcher, E., Farias, S. T., DeCarli, C., & Reed, B. (2018). Education amplifies brain atrophy effect on cognitive decline: implications for cognitive reserve. Neurobiology of Aging, 68, 142–150. doi:10.1016/j.neurobiolaging.2018.04.002

Natale, B. N., Manuck, S. B., Shaw, D. S., Matthews, K. A., Muldoon, M. F., Wright, A. G. C., & Marsland, A. L. (2023). Systemic Inflammation Contributes to the Association Between Childhood Socioeconomic Disadvantage and Midlife Cardiometabolic Risk. Annals of Behavioral Medicine, 57(1), 26–37. doi:10.1093/abm/kaac004

Newton, P., Tchounguen, J., Pettigrew, C., Lim, C., Lin, Z., Lu, H., . . . Team, B. R. (2023). Regional White Matter Hyperintensities and Alzheimer’s Disease Biomarkers Among Older Adults with Normal Cognition and Mild Cognitive Impairment. Journal of Alzheimer’s Disease, 92(1), 323–339. doi:10.3233/JAD-220846

Nielsen, L., Marsland, A. L., Hamlat, E. J., & Epel, E. S. (2024). New Directions in Geroscience: Integrating Social and Behavioral Drivers of Biological Aging. Psychosomatic Medicine, 86(5), 360–365. doi:10.1097/PSY.0000000000001320

Pohl, D. J., Seblova, D., Avila, J. F., Dorsman, K. A., Kulick, E. R., Casey, J. A., & Manly, J. (2021). Relationship between Residential Segregation, Later-Life Cognition, and Incident Dementia across Race/Ethnicity. International Journal of Environmental Research and Public Health, 18(21). doi:10.3390/ijerph182111233

Powell, W. R., Buckingham, W. R., Larson, J. L., Vilen, L., Yu, M., Salamat, M. S., . . . Kind, A. J. H. (2020). Association of Neighborhood-Level Disadvantage With Alzheimer Disease Neuropathology. JAMA Netw Open, 3(6), e207559. doi:10.1001/jamanetworkopen.2020.7559

Prins, N. D., & Scheltens, P. (2015). White matter hyperintensities, cognitive impairment and dementia: an update. Nature Reviews: Neurology, 11(3), 157–165. doi:10.1038/nrneurol.2015.10

Rodriguez, F. S., Lampe, L., Gaebler, M., Beyer, F., Baber, R., Burkhardt, R., . . . Witte, A. V. (2023). Differences in white matter hyperintensities in socioeconomically deprived groups: results of the population-based LIFE Adult Study. International Psychogeriatrics, 1-14. doi:10.1017/S104161022300025X

Rosso, A. L., Flatt, J. D., Carlson, M. C., Lovasi, G. S., Rosano, C., Brown, A. F., . . . Gianaros, P. J. (2016). Neighborhood Socioeconomic Status and Cognitive Function in Late Life. American Journal of Epidemiology, 183(12), 1088–1097. doi:10.1093/aje/kwv337

Sajewski, A. N., Santini, T., DeFranco, A., Berardinelli, J., Jin, H., Li, J., . . . Ibrahim, T. S. (2025). RF shimming strategy for an open 60-channel RF transmit 7T MRI head coil for routine use on the single transmit mode. Magn Reson Med. doi:10.1002/mrm.30563

Santini, T., Kim, J., Wood, S., Krishnamurthy, N., Farhat, N., Maciel, C., . . . Ibrahim, T. S. (2018). A new RF transmit coil for foot and ankle imaging at 7T MRI. Magn Reson Imaging, 45, 1–6. doi:10.1016/j.mri.2017.09.005

Santini, T., Wood, S., Krishnamurthy, N., Martins, T., Aizenstein, H. J., & Ibrahim, T. S. (2021). Improved 7 Tesla transmit field homogeneity with reduced electromagnetic power deposition using coupled Tic Tac Toe antennas. Sci Rep, 11(1), 3370. doi:10.1038/s41598-020-79807-9

Schmidt, R., Berghold, A., Jokinen, H., Gouw, A. A., van der Flier, W. M., Barkhof, F., . . . Group, L. S. (2012). White matter lesion progression in LADIS: frequency, clinical effects, and sample size calculations. Stroke, 43(10), 2643–2647. doi:10.1161/STROKEAHA.112.662593

Seo, S. W., Im, K., Lee, J. M., Kim, S. T., Ahn, H. J., Go, S. M., . . . Na, D. L. (2011). Effects of demographic factors on cortical thickness in Alzheimer’s disease. Neurobiology of Aging, 32(2), 200–209. doi:10.1016/j.neurobiolaging.2009.02.004

Shaked, D., Leibel, D. K., Katzel, L. I., Davatzikos, C., Gullapalli, R. P., Seliger, S. L., . . . Waldstein, S. R. (2019). Disparities in Diffuse Cortical White Matter Integrity Between Socioeconomic Groups. Frontiers in Human Neuroscience, 13, 198. doi:10.3389/fnhum.2019.00198

Sheets, L. R., Henderson Kelley, L. E., Scheitler-Ring, K., Petroski, G. F., Barnett, Y., Barnett, C., . . . Parker, J. C. (2020). An index of geospatial disadvantage predicts both obesity and unmeasured body weight. Prev Med Rep, 18, 101067. doi:10.1016/j.pmedr.2020.101067

Sims, M., Kershaw, K. N., Breathett, K., Jackson, E. A., Lewis, L. M., Mujahid, M. S., . . . Outcomes, R. (2020). Importance of Housing and Cardiovascular Health and Well-Being: A Scientific Statement From the American Heart Association. Circulation: Cardiovascular Quality and Outcomes, 13(8), e000089. doi:10.1161/HCQ.0000000000000089

Tan, C. H., & Tan, J. J. X. (2023). Low neighborhood deprivation buffers against hippocampal neurodegeneration, white matter hyperintensities, and poorer cognition. Geroscience, 45(3), 2027–2036. doi:10.1007/s11357-023-00780-y

Thomas, A. J., Hamilton, C. A., Donaghy, P. C., Martin-Ruiz, C., Morris, C. M., Barnett, N., . . . O’Brien, J. T. (2020). Prospective longitudinal evaluation of cytokines in mild cognitive impairment due to AD and Lewy body disease. International Journal of Geriatric Psychiatry, 35(10), 1250–1259. doi:10.1002/gps.5365

Tosto, G., Zimmerman, M. E., Carmichael, O. T., Brickman, A. M., & Alzheimer’s Disease Neuroimaging, I. (2014). Predicting aggressive decline in mild cognitive impairment: the importance of white matter hyperintensities. JAMA Neurol, 71(7), 872–877. doi:10.1001/jamaneurol.2014.667

Vassilaki, M., Aakre, J. A., Castillo, A., Chamberlain, A. M., Wilson, P. M., Kremers, W. K., . . . Petersen, R. C. (2022). Association of neighborhood socioeconomic disadvantage and cognitive impairment. Alzheimers Dement. doi:10.1002/alz.12702

Waldstein, S. R., Dore, G. A., Davatzikos, C., Katzel, L. I., Gullapalli, R., Seliger, S. L., . . . Zonderman, A. B. (2017). Differential Associations of Socioeconomic Status With Global Brain Volumes and White Matter Lesions in African American and White Adults: the HANDLS SCAN Study. Psychosomatic Medicine, 79(3), 327–335. doi:10.1097/PSY.0000000000000408

Walhovd, K. B., Fjell, A. M., Wang, Y., Amlien, I. K., Mowinckel, A. M., Lindenberger, U., . . . Brandmaier, A. M. (2022). Education and Income Show Heterogeneous Relationships to Lifespan Brain and Cognitive Differences Across European and US Cohorts. Cerebral Cortex, 32(4), 839–854. doi:10.1093/cercor/bhab248

Williams, D. R., Mohammed, S. A., Leavell, J., & Collins, C. (2012). Race, socioeconomic status, and health: Complexities, ongoing challenges, and research opportunities. Annals of the New York Academy of Sciences, 1186, 69–101.

Xu, J., Lawrence, K. G., O’Brien, K. M., Jackson, C. L., & Sandler, D. P. (2022). Association between neighbourhood deprivation and hypertension in a US-wide Cohort. Journal of Epidemiology and Community Health, 76(3), 268–273. doi:10.1136/jech-2021-216445

Zuelsdorff, M., Larson, J. L., Hunt, J. F. V., Kim, A. J., Koscik, R. L., Buckingham, W. R., . . . Kind, A. J. H. (2020). The Area Deprivation Index: A novel tool for harmonizable risk assessment in Alzheimer’s disease research. Alzheimers Dement (N Y), 6(1), e12039. doi:10.1002/trc2.12039

